# A novel method for tri-clustering dynamic functional network connectivity (dFNC) identifies significant schizophrenia effects across multiple states in distinct subgroups of individuals

**DOI:** 10.1101/2020.08.06.239152

**Authors:** Md Abdur Rahaman, Eswar Damaraju, Jessica A. Turner, Theo G.M. van Erp, Daniel Mathalon, Jatin Vaidya, Bryon Muller, Godfrey Pearlson, Vince D. Calhoun

## Abstract

**Background:** Brain imaging data collected from individuals are highly complex with unique variation; however, such variation is typically ignored in approaches that focus on group averages or even supervised prediction. State-of-the-art methods for analyzing dynamic functional network connectivity (dFNC) subdivide the entire time course into several (possibly overlapping) connectivity states (i.e., sliding window clusters). Though, such an approach does not factor in the homogeneity of underlying data and may end up with a less meaningful subgrouping of the dataset.

**Methods:** Dynamic-N-way tri-clustering (dNTiC) incorporates a homogeneity benchmark to approximate clusters that provide a more apples-to-apples comparison between groups within analogous subsets of time-space and subjects. dNTiC sorts the dFNC states by maximizing similarity across individuals and minimizing variance among the pairs of components within a state.

**Results:** Resulting tri-clusters show significant differences between schizophrenia (SZ) and healthy control (HC) in distinct brain regions. Compared to HC, SZ in most tri-clusters show hypoconnectivity (low positive) among subcortical, default mode, cognitive control but hyper-connectivity (high positive) between sensory networks. In tri-cluster 3, HC subjects show significantly stronger connectivity among sensory networks and anticorrelation between subcortical and sensory networks compared to SZ. Results also provide statistically significant difference in reoccurrence time between SZ and HC subjects for two distinct dFNC states.

**Conclusions:** Outcomes emphasize the utility of the proposed method for characterizing and leveraging variance within high-dimensional data to enhance the interpretability and sensitivity of measurements in the study of a heterogeneous disorder like schizophrenia and in unconstrained experimental conditions such as resting fMRI.

## Introduction

Heterogeneity in schizophrenia represents a challenge for the study and diagnosis of this disorder (Alnæs et al., 2019; Rahaman et al., 2019; Tsuang, Lyons, & Faraone, 1990). A substantial amount of research effort has focused on identifying the causes of schizophrenia, that would aid in improving diagnosis and treatments, yet we are still not close to the root cause of this mental illness (Du et al., 2019; Ferri et al., 2018; Lawrie & Abukmeil, 1998; B. Rashid et al., 2019). Studies often use structural magnetic resonance imaging (sMRI), resting-state functional MRI (fMRI), and task-based fMRI to analyze a wide range of neurocognitive variables; for instance, brain network activation, subtyping, neural components clustering, etc. (E. A. Allen et al., 2011; V. D. Calhoun, Kiehl, & Pearlson, 2008; Rahaman et al., 2019). Several research efforts have made progress on studying brain connectivity using structural imaging data and task-based MRI analysis (Arbabshirani, Havlicek, Kiehl, Pearlson, & Calhoun, 2013; Clayden, 2013; Di et al., 2015; Gonzalez-Castillo & Bandettini, 2018). Task-related functional connectivity (FC) studies of schizophrenia have reported reliable FC differences across several visual regions and suggested task-based dynamic methods could improve our understanding of the coordination between different activated brain networks (Di et al., 2015). Resting-state fMRI is a widely used method for exploring neural activity - because it has the benefit of being both easy for patients to perform and potentially more sensitive to brain disorders (Damoiseaux et al., 2006; Franco, Pritchard, Calhoun, & Mayer, 2009; Shehzad et al., 2009; Zuo et al., 2010). Studies during rest fMRI have identified several temporally coherent networks that are putatively involved in functions such as vision, audition, and directing attention (Beckmann, DeLuca, Devlin, & Smith, 2005; V. D. Calhoun, Adali, Pearlson, & Pekar, 2001). More recent studies have exploited the dynamics and intrinsic fluctuations of connectivity, a property that has been exploited in various studies (Arieli, Sterkin, Grinvald, & Aertsen, 1996; Kucyi & Davis, 2014; Leonardi & Van De Ville, 2015; Makeig, Debener, Onton, & Delorme, 2004; Onton, Westerfield, Townsend, & Makeig, 2006).

Dynamic functional network connectivity (dFNC) is a promising analysis approach that provides time-varying correlation matrices typically computed using a sliding window (e.g. 44s in length) technique for calculating the correlations among brain regions or independent components of interest (Elena A Allen et al., 2014; Damaraju et al., 2014; Hindriks et al., 2016; Ioannides, 2007). Here, each correlation can be considered as a transient functional association between a pair of functional networks (Ioannides, 2007). These study of temporal variations in functional connectivity (FC) among spatially independent neural systems have already been established as a broader avenue for studying neurodegenerative diseases (Fu et al., 2018; Klugah-Brown et al., 2019; Ma, Calhoun, Phlypo, & Adalı, 2014; Vergara, Mayer, Kiehl, & Calhoun, 2018; Wang et al., 2019). Studies implement dFNC for improving classification accuracy (Cetin et al., 2016; Barnaly Rashid et al., 2016; Sakoglu, Michael, & Calhoun, 2009) and some studies found discontinuous connectivity by experimenting with connectivity states (Du et al., 2017). Functional connectivity patterns in SZ are considered to be informative of this disorder and some reportedly popular neural features in the literature are the reduced amplitude of low-frequency fluctuation (ALFF) in cuneus (Hoptman et al., 2010; J. A. Turner et al., 2013) and dysconnectivity between thalamus and sensory regions (Vince D Calhoun, Eichele, & Pearlson, 2009; Kühn & Gallinat, 2013; Malaspina et al., 2004; Mingoia et al., 2012; Zhou et al., 2007). Moreover, researchers have focused on the connectivity patterns of dFNC in SZ that provide crucial insights about the brain dynamics which might not be available in static investigations (Ma et al., 2014; Barnaly Rashid et al., 2016). However, in the previous studies, the analysis performed on dFNC windows combines all individuals for capturing the groupings, connectivity patterns and changes in brain networks over time (Elena A Allen et al., 2014; Damaraju et al., 2014; Miller et al., 2016a; Shakil, Lee, & Keilholz, 2016). These studies partition the whole dynamic into a certain number of states based on the homogeneity in the connectivity pattern using a clustering technique. Essentially, the technique assigns all the dFNC windows into states but does not directly optimize for their subject-wise consistency. Secondly, the windows include all the considering functional networks which implies the connectivity patterns are spread throughout the neural system in these studies. For the connectivity signatures that are more transient and has a comparatively lesser scope i.e., beyond a non-contiguous smaller subset of brain functional units, the previous methods would fail to capture the patterns and reflect them in clustering process. The clustering approach rather average over the shorter motifs to build a more general form of connectivity. Consequently, it stringent our capability to investigate the differences between distinct group of subjects. Here, we argued the necessity of further sorting the subset of subjects, pair of components, and time points enclosed within the dFNC states. So, we propose to cluster all three dimensions (subjects, windows, and network pairs) of the dataset together for extracting more homogenous subgroups to provide better comparison.

In three-dimensional data, observations are generally meaningfully correlated across various dimensions of the dynamic. As such, one-dimensional clustering is not likely to provide the most effective mean of exploration in this scenario. Although biclustering takes care of this challenge partially and has been shown to be more effective, it does not include the third dimension (Rahaman et al., 2019). As dFNC is a complex three-dimensional representation of the data, by not clustering all three dimensions we might be missing significant group differences in the data. dNTiC simultaneously clusters three neural variables and return subgroups of entities with a maximized homogeneity across those given variables. Our approach, to our knowledge, represents the first attempt at exploring all three dimensions of dFNC as well as a robust method of tri-clustering multiple neural features. We perform more rigorous sorting on the subject within a state while keeping each of their windows to be highly correlated with the cluster centroid. dNTiC compact the connectivity patterns by ignoring the heterogeneous connections and weakly correlated subjects; results show it still identifies more group differences than the one-dimensional k-means approach, presumably because comparisons are more stable and meaningful (Damaraju et al., 2014). These tri-clusters report SZ-HC group differences in figure 5. In earlier study, the states were very similar for both SZ and HC group, therefore, the differences are very subtle. Here, we observe SZ < HC connectivity strength across Auditory (AUD), Visual (VIS), and Sensorimotor (SM) regions and SZ > HC at subcortical (SC) in tri-clusters 3 and 5 (fig 4). In tri-cluster 1, we observe SZ > HC in AUD, VIS, and SM domains. The experiment on reoccurrence time reveals two subgroups showing statistically significant difference between patient and control (tri-cluster 2 and 5). Overall, HC subjects show higher reoccurrence time than SZ except subgroup 2.

## Data collection and preprocessing

In this study, we used resting-state fMRI data collected as part of the Functional Imaging Biomedical Informatics Research Networks (fBIRN) project consisting of 163 healthy controls and 151 age-and gender-matched patients with schizophrenia in a total of 314 subjects (Keator et al., 2016; Jessica A Turner et al., 2013). The dataset includes 117 males, 49 females; mean age of 36.9 from healthy cohort and 114 males, 37 females; mean age of 37.8 from patients with schizophrenia. More demographics of the samples are available in the above references and also reported in (Damaraju et al., 2014). The data acquisition and preprocessing follow the similar pipeline as described in Damaraju et al., 2014 (Damaraju et al., 2014). In addition, data quality control, head motion correction, diagnosis, etc. is also described in the supplementary document of this study. The collecting study ensured the informed consent of all participants before scanning. Briefly, a blood oxygenation level-dependent (BOLD) fMRI scan was performed on 3T scanners across 7 sites with patients and controls collected at all sites. Resting-state fMRI scans were acquired using the parameters FOV of 220 × 220 mm (64 × 64 matrix), TR = 2 s, TE = 30 ms, FA = 770, 162 volumes, axial slices = 32, slice thickness = 4 mm and skip = 1 mm. The scan was a closed eye resting-state fMRI acquisition. Image preprocessing was conducted using several toolboxes namely AFNI^1^, SPM^2^, GIFT^3^, and custom MATLAB scripts. First, rigid body motion correction was performed using SPM to correct for subject head motion and slice-timing correction to account for timing differences in slice acquisition. Then time series data is despiked to mitigate the outlier effect and warped to a Montreal Neurological Institute (MNI) template and resampled to 3 mm^3^ isotropic voxels and followed by smoothing and variance normalization. We smoothed the data to 6 mm full width at half maximum (FWHM) using AFNI’s BlurToFWHM algorithm. After finishing the preprocessing, we ran a group independent component analysis (GICA) (V. D. Calhoun et al., 2001; Erhardt et al., 2011) on the functional data. We performed group ICA on the preprocessed data and identified 47 intrinsic connectivity networks (ICNs) from the decomposition of 100 components. Spatial maps (SMs) and time courses (TCs) for each subject were obtained using the spatiotemporal regression back reconstruction approach (V. D. Calhoun et al., 2001). Subject wise SMs and TCs are then post-processed as described in Allen et al., 2012 (Elena A Allen et al., 2014). The dynamic FNC (dFNC) between two ICA time courses was computed using a sliding window approach with a window size of 22 TR (44 s) in steps of 1 TR. The sliding window is a rectangular window of 22 time points convolved with Gaussian of sigma 3 TRs to obtain tapering along the edges (Elena A Allen et al., 2014). We computed covariance from regularized inverse covariance matrix (ICOV) (Smith et al., 2011; Varoquaux, Gramfort, Poline, & Thirion, 2010) using graphical Lasso framework (Friedman, Hastie, & Tibshirani, 2008). Moreover, to ensure sparsity, we imposed an additional L1 norm constraint on the inverse covariance matrix. We used a log-likelihood of unseen data of the subject in a cross-validation framework for optimizing regularization parameters. To ensure positive semi-definiteness of the evaluated dynamic covariance matrices, we ensure their estimated eigen values are positive. Then we compute the dFNC values for each subject. And the covariance values are Fisher-Z transformed and residualized with respect to age, gender, and sites.

## Methods

Our methodology consists of three fundamental steps, 1) generating dFNC states, 2) sorting states, and 3) exploring subsets of states as shown in figure 3. Figure 1 shows the different dimensions of our dFNC dataset where the dFNC time course of each subject has a length of 136 windows and each window consist of 1081 pairs – each pair represents a correlation value between two independent brain networks scaled from −1 to 1. From one hundred ICA components, 47 were reported as relevant intrinsic brain networks in the earlier study which generates a 47 × 47 symmetric matrix of connectivity (Damaraju et al., 2014). Thus, by taking either part of the diagonal we obtain a vector of component’s pair of size 1081 (47 choose 2).

To partition the dynamics into dFNC states, we collect windows from all subjects of the dataset and run *k*-means clustering based on geometric distance with an optimal k. For selecting the model order (k), we used the elbow criteria, the ratio of within-cluster to between cluster distances as suggested in previous studies (Abrol et al., 2017; Miller et al., 2016b; D. K. Saha et al., 2019; Debbrata K. Saha et al., 2020). We observe elbow for k = 5 (Fig 2). We describe these window clusters as ‘k’ different dFNC states. Theoretically, windows within a state are expected to be analogous in terms of connectivity patterns. A state is consisting of a subset of subjects having at least one window included in that cluster. So, a subject can be included in the maximum k number of states and a minimum in 1 state.

The second step is sorting the states estimated from k-means for subjects and pairs. As described above, each state is a set of windows and we can infer a subset of subjects by assigning their membership based on the fact that, each of those subjects has at least one window or a set of windows highly correlated to the centroid of corresponding state/cluster. Similarly, selecting those pairs which show low variance across the subjects included in a state. We described both sorting techniques elaborately in the following paragraphs.

### Sorting subjects within a state (maximizing the correlation)

Here we raise the bar of uniformity and filter out the samples showing erratic patterns by maximizing homogeneity across the windows within a cluster. For each subject with a state k, we compute a mean window over all the windows of that subject included in state k and we describe it as the mean window (μW) of type k. Next, the method checks the similarity between the subject wise mean window (μW) and the centroid of a particular state C_k_ based on the Spearman correlation between them. It quantifies that correlation for all the subjects within a state and evaluates the average which is defined as a mean association (μα) by all the subjects per state. Finally, for assigning membership to the subject, the process uses the following equations which retain subjects which have a higher association with the cluster centroid than a mean association. A subject is included in state S if,

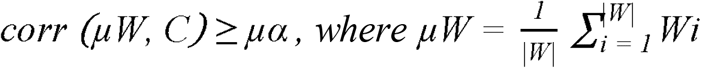

where |W| is the number of windows of the subject included in state k, C is the centroid of a cluster and ‘corr’ is a MATLAB function for computing correlation between two vectors. At the end of this process, the algorithm generates a subset of strongly connected subjects for each of the dFNC states.

### Sorting the pairs within a state (minimizing the variance)

Here, the approach optimizes a subset of network pairs that show similar patterns of connectivity in all the windows, which is basically in all the subjects of a state k. Windows of type k from all the sorted subjects are included in a state to create a collection of windows and then apply variance minimization on each window across the subjects. For each pair, the algorithm computes variance over all the windows (σ^2^) and every window has 1081 pairs which yield 1081 variances. Then, it evaluates the mean of variances (μσ^2^). Our approach sorts each state for a subset of pairs showing relatively lower variance across the sorted subjects. We essentially select pairs with variance < mean of variances. A pair p is included in state S if,

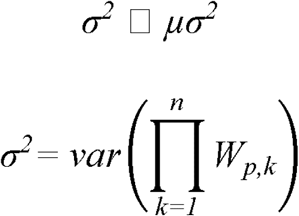

where W_p,k_ represents p^th^ value in k^th^ window and n correspond to the number of windows in the state S.

### mDFS subroutine

The third step is exploring all possible subsets of sorted dFNC states. The algorithm uses a modified depth-first search (mDFS) (Even, 2011) for generating all possible subsets of states. The intuition is to explore the homogeneity across multiple states. Since we have five dFNC states, the analysis assigns the minimum number of states parameter from 1 to 5. However, dFNC state is a three-dimensional substructure of subjects, pairs, and windows thus, fusing more states into a tri-cluster quickly leads to a tiny shared subspace since we intersect across the features (subjects, pairs, windows) of considering states. Therefore, it is cardinal to set the appropriate input parameters for extracting meaningful subgroups and parameters required to specify before running dNTiC algorithms are as follows.

S: List of sorted states (ex: 1, 2, 3, 4, 5, etc.)

N: Minimum number of subjects in a tri-cluster

M: Minimum number of states/window cluster in a tri-cluster

P: Minimum number of pairs in a tri-cluster

O: Allowed percentage of overlap between two tri-clusters

For optimizing the parameters, first, we specify ‘O’ – allowed percentage of overlap between the tri-clusters. We explore multiple values for that parameter which controls overlap among extracted dTiCs. We evaluate percentages from 5 to 35 with an interval of 5. This is a selection user can tweak around to perturb the analysis and the choice is vastly dependent on the dataset, number of subjects included in the sorted states, and the average overlap in subjects between the states (after k-means). However, we can get an idea of these free selections from the remaining subjects, pairs, and windows in the states we are analyzing. Table 1 shows the subgroups before and after sorting the states according to step 1 and 2. In our case, state 3 has the minimum number of subjects (S = 73), so, intersection with this state can yield maximum 73 subjects in a dTiC if all the subjects match with other comparing state. However, for generalization, after multiple runs, we set them to moderate number e.g., N = 50, M = 1 or 2 and P = 500. We present results on dTiCs that include a single state since the sorting steps already downsize the feature’s collection to a significant degree. So, these tri-clusters can provide imperative results on connectivity signatures and group differences. The parameters selection and use cases are partially explained in (Rahaman et al., 2020). For creating the subsets, the mDFS uses the parameter *M* representing the minimum number of states in a tri-cluster (dTiC). For restricting exponential growth of the search space, we tweaked the depth-first search (DFS) on the set of states. For in backtracking from a branch after identifying an arbitrary sized (*m*) subsets, it checks the boundary conditions for N, M, and O and any violation leads to the earlier branch of search tree. For more clarity, we consider the states serial numbers (1, 2, 3, 4, 5) as the nodes from an undirected graph and aim to establish edges (subset) between them based on satisfactory conditions on the input parameters. An unsatisfied condition might be the number of subjects are less than or the overlap becomes greater than the threshold calculated based on *O*. Early abandoning a branch is possible (backtracking) in our case because the major operation in the searching process is intersection. Theoretically, if the earlier intersection results in a tri-cluster (dTiC) with an inadequate number of subjects/pairs/windows then there is no chance of obtaining a valid subgroup in further steps. As such, the algorithm stops exploring the path and backtracks to an earlier point. For each subset of states, the intersection is performed in both subject and pair dimensions. So, for each eligible subset of states, we obtain an intersected 3D subspace consisting of a subset of states, subjects, and pairs. After that, this subspace (3D submatrix) scrutinized for overlapping parameter by checking the ratio of overlap with the tri-clusters that have already been listed. Here we use the F_1_ similarity index to investigate the overlap between any earlier reported dTiCs (Hripcsak & Rothschild, 2005; Santamaria, Quintales, & Therón, 2007). The F_1_ similarity index is defined as follows for any two arbitrary tri-clusters A and B,

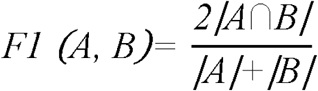

|A∩B| =Intersection between dTiC A and B; S_A∩B_ × W_A∩B_ × P_A∩B_

|A|= Size of A; i.e. the number of subjects × windows × pairs

|B| = Size of B; i.e. the number of subjects × windows × pairs

Based on positive feedback from the validator, our method adds the dTiC into a list of tri-clusters. After traversing through the whole search space by mDFS, dNTiC returns the list of tri-clusters as the final output of the investigation.

**Table 1.**
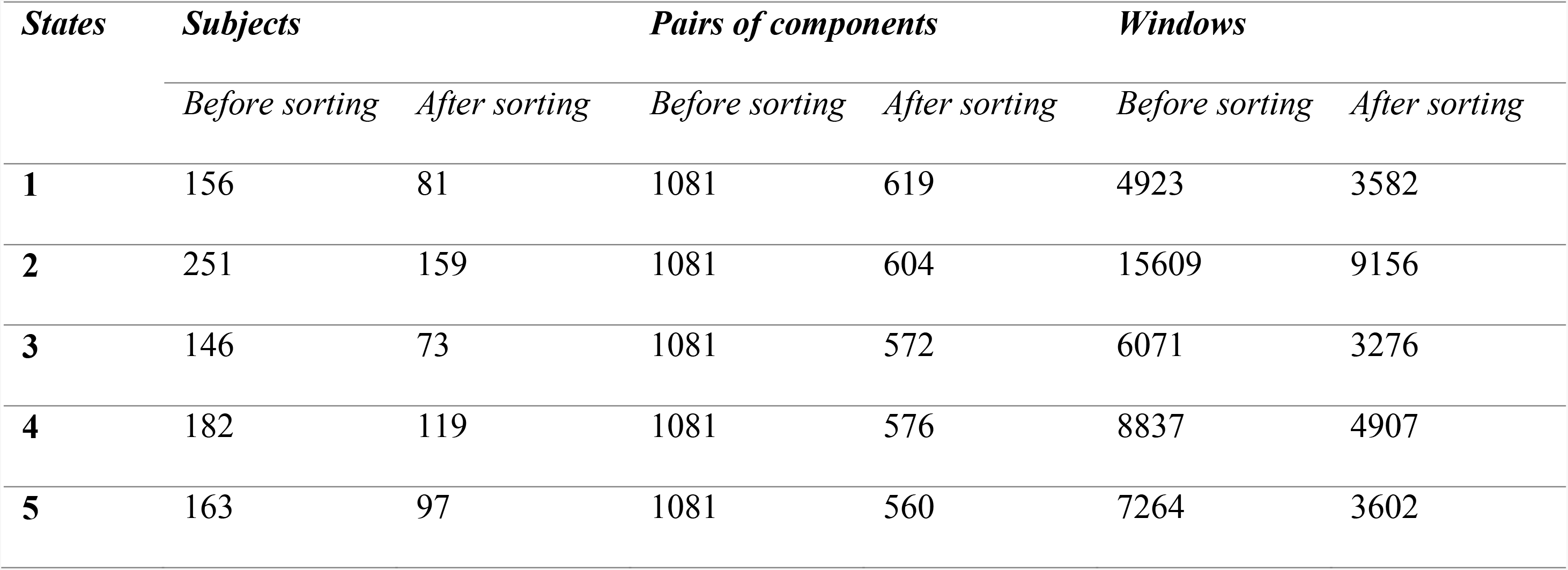
Characteristics of the sorted dFNC states. The table demonstrates number of subjects, pairs of components, and windows included in the dFNC states before and after running sorting process (step1 and 2 of our methodology)

## Results

We run the experiments using distinct combinations of input parameters for dNTiC approach. The significant results are presented in the following section also, provide results from the extended investigation in our supplementary documents. Our method extracts 5 tri-clusters (dTiC) for the input argument N = 50, M =1 and P = 500. In figure 4, we present the group differences identified from these tri-clusters by running a two-sample t-test on each pair of components. In two upper rows, most of the dTiCs show higher connectivity in subcortical (SC), visual (VIS), sensorimotor (SM), and default mode (DM) domains. The SC components show mild to moderate anticorrelation with auditory (AUD), VIS, SM regions across the states and low negative to positive connectivity to all other.

dTiC 3 is the subgroup that shows strong group differences among all the estimated dynamic tri-clusters. dTiC 1 and 3 distinguish each other in DMN connectivity (anti-correlated strongly to VIS and Motor in 1 vs 3). dTiC 3 and 5 differ in SC connectivity to other domains and dTiC 2 and 4 also distinguish each other in DMN connectivity. Apparently, in dTiC 1, 3 and 5 have more active pairs in VIS and SM regions than dTiC 2 and 4. However, in dTiC 2 and 4 have lightly more pairs in the cognitive control (CC) region. These are distinct tri-clusters of subjects and pairs where one group of tri-clusters (dTiC 1, 3, and 5) is more active/dense across different regions whereas, another group (dTiC 2 and 4) is sparsely active which means fewer pairs clustered together in those subjects.

The bottom row (of fig 4) demonstrates the group differences reported by the dTiCs. In dTiC 1, we can see pairs from SZ individuals showing higher connectivity strength than HC pairs in VIS and SM regions. dTiC 2 is mostly much sparse in different regions except SZ-HC hyper-connectivity in DMN and CC. In contrast, dTiC 3 shows group differences across several regions of the brain. Here, healthy control (HC) showing a stronger within domain connectivity in visual (VIS), auditory (AUD) and sensorimotor (SM) regions and greater anti-correlation to SC and CC nodes comparing to SZ. The directionality in the VIS-SM region is HC > SZ and others are reverse. We find strong group differences in subcortical to AUD-VIS-SM domains and it is higher correlations in the SZ group and lower in HC. Patients exhibit reduced connectivity (in this case lack of anti-correlation) compared to HC. We also observe group differences in functional connectivity from DM to all other domains that evident a considerable effect of SZ in the subject’s neural system. dTiC 4 shows significant differences in DM-VIS, CC and dTiC 5 shows group difference in connectivity in SC to all other domains. The last dTiC is similar to dTiC 3 to some extent unless both of them distinguish each other in cognitive control components. In essence, subcortical, sensory, default mode, and cognitive control regions revealed significant differences in average connectivity strength between patients and controls. In an analysis where we forced dNTiC to explore homogeneity across more than two states, we observed a large and significant tri-cluster (dTiC 4) where states 4 and 5 were clustered together and show significant group differences in sensory and default mode regions. We explain this scenario in the supplement.

Now, we evaluated the transition of each subject throughout these three-dimensional homogenous dynamic patterns. We measure window frequency for each subject which represents how frequently a subject change its pattern also, how long it continues with the same pattern. In fig 5, we computed window frequency of each subject within a tri-cluster which represents the percentage of windows type ‘i’ for that subject. A subject can be a member of multiple tri-clusters. For a subject taken from dTiC ‘i’ we compute the percentage of the window of type ‘i’ which is also the amount of time in TR the subject spends in state ‘i’. These negative t-values are evident that window frequency in SZ subjects is smaller than in HC. Thus, HC subjects linger within uniform connectivity states whereas SZ subjects move back and forth among relatively shorter states. However, of dFNC time course, SZ subjects in dTiC 2 have more windows of type 2 which evidence these subjects’ dynamics spent more time in state 2 than the HC subjects within. Overall, the results demonstrate that a quicker transition among different connectivity states better characterize SZ subjects.

We check the correlation between mean functional connectivity and the positive and negative syndrome scale (PANSS) score from the SZ subjects to investigate if these patterns are related to the symptoms of the disorder (Kay, Fiszbein, & Opler, 1987). The regression line characterizes the pattern of symptom scores (positive, negative, and general) with the change of average connectivity strength of the patients included in a subgroup. We see from fig 6, dTiC 1 anticorrelated with positive and associated with negative symptoms; subjects in dTiC 2 express bold general symptoms and the corresponding regression line demonstrates linear increment with the rise of connectivity. SZ subjects included in dTiC 3 show a unique trend in general symptom scores compared to all other subgroups. The general symptom score decreases with the rise of mean connectivity among different functional networks of the brain, on the other hand, it increases with the connectivity strength and significantly correlated with the general symptoms (i.e., anxiety, guilt feelings, tension, etc.). Moreover, the subgroup expresses more positive symptoms with higher connectivity and for the negative symptom, it follows the opposite. SZ subjects in dTiC 4 show a moderate correlation with negative PANSS scores and the symptoms are expressed more strongly with the increase of connectivity which is a unique pattern within this subgroup. Lastly, the dTiC 5 also exhibits association with positive symptoms which manifest with higher connectivity. Although, the associations are not very strong in term of the correlation value, but these indicate a few trends depicting interactions between the symptom scores and the connectivity strength within the subgroups.

In figure 7, we present the group differences in reoccurrence time of each state between patients and control. Reoccurrence time of a tri-cluster is given by the number of windows assigned to that dTiC. So, from each subject’s windows how many are assigned to a tri-cluster is the reoccurrence time for that subject. The definition is consistent with earlier study (Li et al., 2017). A two-sample t-tests reveal dTiC 2 and 5 showing statistically significant (at a level *p* < 0.05) group differences in the reoccurrence time. We observe SZ subjects show a higher reoccurrence time for dTiC 3 but lower reoccurrence time in dTiC 5. It indicates the connectivity pattern depicted by dTiC 2 is comparatively more recurring in SZ subjects than any other connectivity patterns.

## Discussion

The study demonstrates a novel method of tri-clustering dFNC that sifts through all three dimensions of the data (network pairs, windows, subjects) simultaneously. dNTiC provides a mean of subgrouping the data more precisely which can reveal significant complex relationships within the subgroups that do not exist across the dimensions. The outcomes of the study are intriguing because throughout the method we maximized the homogeneity across the elements within a tri-cluster and then we analyzed those highly homogenous subspaces for patient-control group differences which makes the reported outcomes of the comparison more sensitive and meaningful. The results reveal significant differences in connectivity, coactivations, antagonism, etc. across a set of distinct brain regions. Subjects from tri-cluster 3 report distinguished group differences at distinct regions of the brain: a sharp antagonism in visual and sensorimotor regions and less anticorrelation between subcortical and sensory networks. This subgroups also shows a higher correlation with positive symptom and anticorrelation with negative and general symptom scores which indicates patient with hypo connectivity in sensory (AUD, VIS, SM) and hyper-connectivity between subcortical and sensory networks are more likely to express positive symptoms (i.e. hallucinations, delusions, racing thoughts. The previous studies appear to reveal comparatively less functional connectivity differences within the brain regions (Barber, Lindquist, DeRosse, & Karlsgodt, 2018; Damaraju et al., 2014; Du et al., 2017; Barnaly Rashid, Damaraju, Pearlson, & Calhoun, 2014). A potential reason for less differences could be the scattered focus throughout the brain. The main contributions of our study are building more comparable subspace by reducing the heterogeneity across the characteristics of dFNC states. That finer comparability helps extracting more group differences than earlier analysis. For a quantitative comparison, we can consider group differences in fig 3(B) of (Damaraju et al., 2014) which used the same dataset and followed an identical preprocessing pipelines. Apparently, our dTiCs provide more SZ - HC group differences in number of domains, where their states report almost no differences except a very sparse dots in AUD, VIS, SM regions. For the reference, we are providing the figure 3 (B) from that study in our supplementary document (see supplementary figure 3). In our study, the tri-clusters are more precisely sorted for a subset of subjects showing minimum heterogeneity in their connectivity pattern and show significant group differences across multiple brain regions. From fig 4, we can see that all the dTiCs includes a subset of subjects which comprises one-third of the total number of subjects (N=314) and half of the total number of pairs (N=1081). This type of subgrouping of subjects and pairs can illuminate domain-wise connectivity differences within patients and control which would be lost in a study using standard *k*-means or any other conventional clustering approaches (Elena A Allen et al., 2014; Damaraju et al., 2014; Miller et al., 2016a; Barnaly Rashid et al., 2014). The findings from the window frequency experiment characterize patients with a higher transition frequency across distinct connectivity states. This sporadic showcase of SZ dynamics also aligns with the previous investigation (Elena A Allen et al., 2014; Damaraju et al., 2014). Further, when we ran the model using an increased number of states in a tri-cluster which is less than equal 2 (see supplement), which highlights the changes in connectivity strength for the extended subgroup that includes multiple states. The intuition behind mounting multiple states in each tri-cluster is to capture longer connectivity patterns which give an approximation over the persistence of the states across the time course. However, the broader tri-clusters report less significant variations in the connectivity pattern rather than replicating the older templates among a smaller subset of connections/pairs of components. We also run the analysis for larger model order (k = 9) where we observe tri-clusters which are consistent to model order 5; indicating starting with more dFNC states eventually converges to the similar connectivity patterns which we are studying here with a lower k

## Conclusion and future directions

Studies of functional connectivity data have largely ignored both individual variabilities and have not optimized across the multiple data dimensions. So, we are looking for clusters in the dataset which may show a relation among the data variables while ensuring the homogeneity within the subgroup. Moreover, it is useful to unveil biomarkers for mental disorders that are heterogeneous, such as schizophrenia and enriches our understanding of how connectivity between components of the brain modulates diverse impact on subjects with this kind of disorder. By calibrating the size parameter of the algorithm, we can also analyze the stability of a certain feature. For instance, if we identify a feature within a subgroup and by increasing the number of states gradually, we can test the longevity of this trend for a greater subset of states. Thus, dNTiC helps not only identifying a feature but also drawing an effect size of this feature. Our framework is largely dependent on the selection of input parameters. We are working on developing an automated framework to select these arguments for dNTiC algorithm based on the given information and other inferences. In the future, we are planning to extend the method for task-based fMRI data in which the windows would be synchronous across the subjects thus we can directly tri-cluster the data without applying k-means and that will potentially increase the robustness of the algorithm. For the sorting steps, we are working on a Gaussian kernel-based tracking technique to identify the coactivation of connectivity patterns across the subjects over the time course.

---

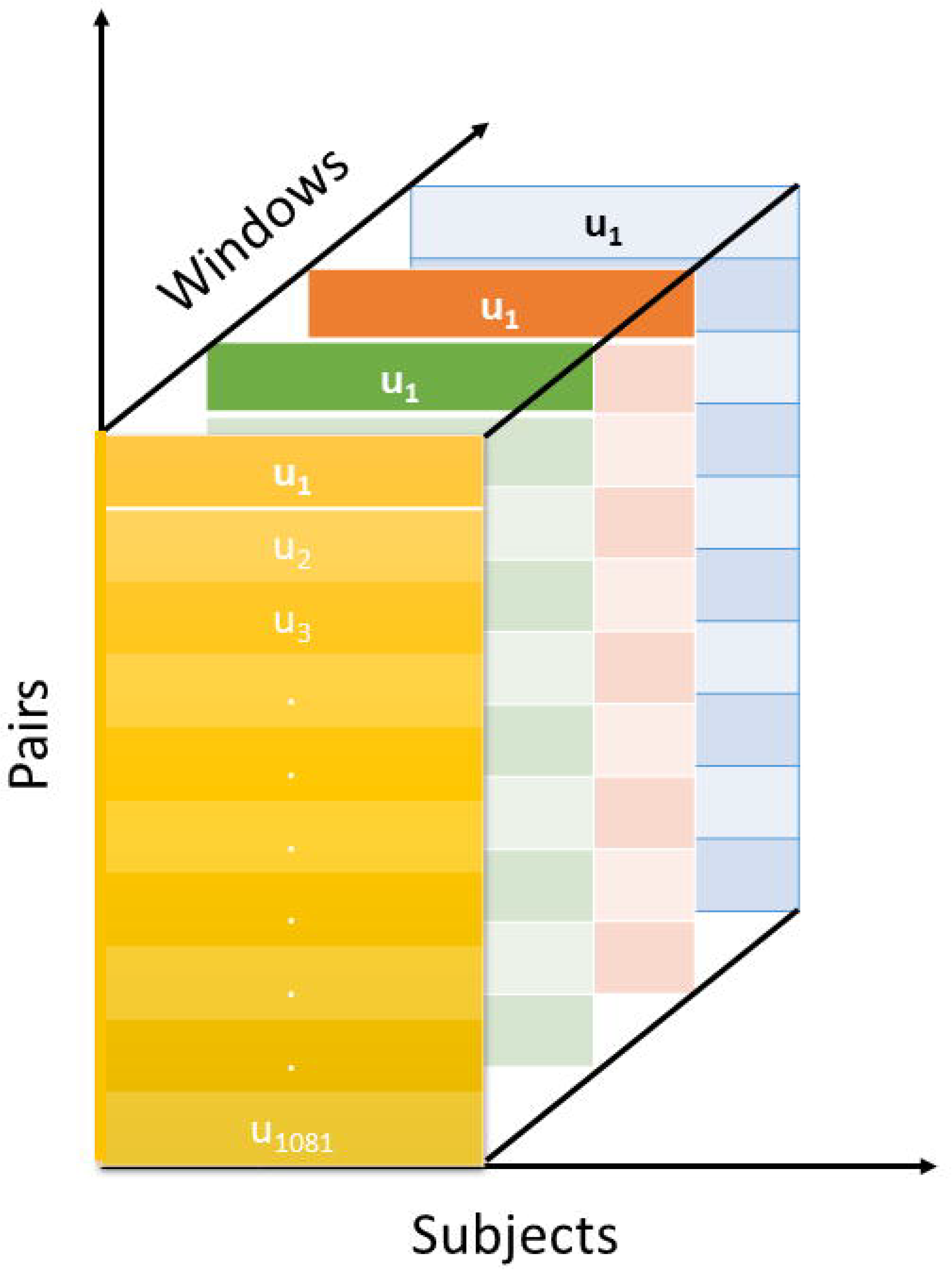

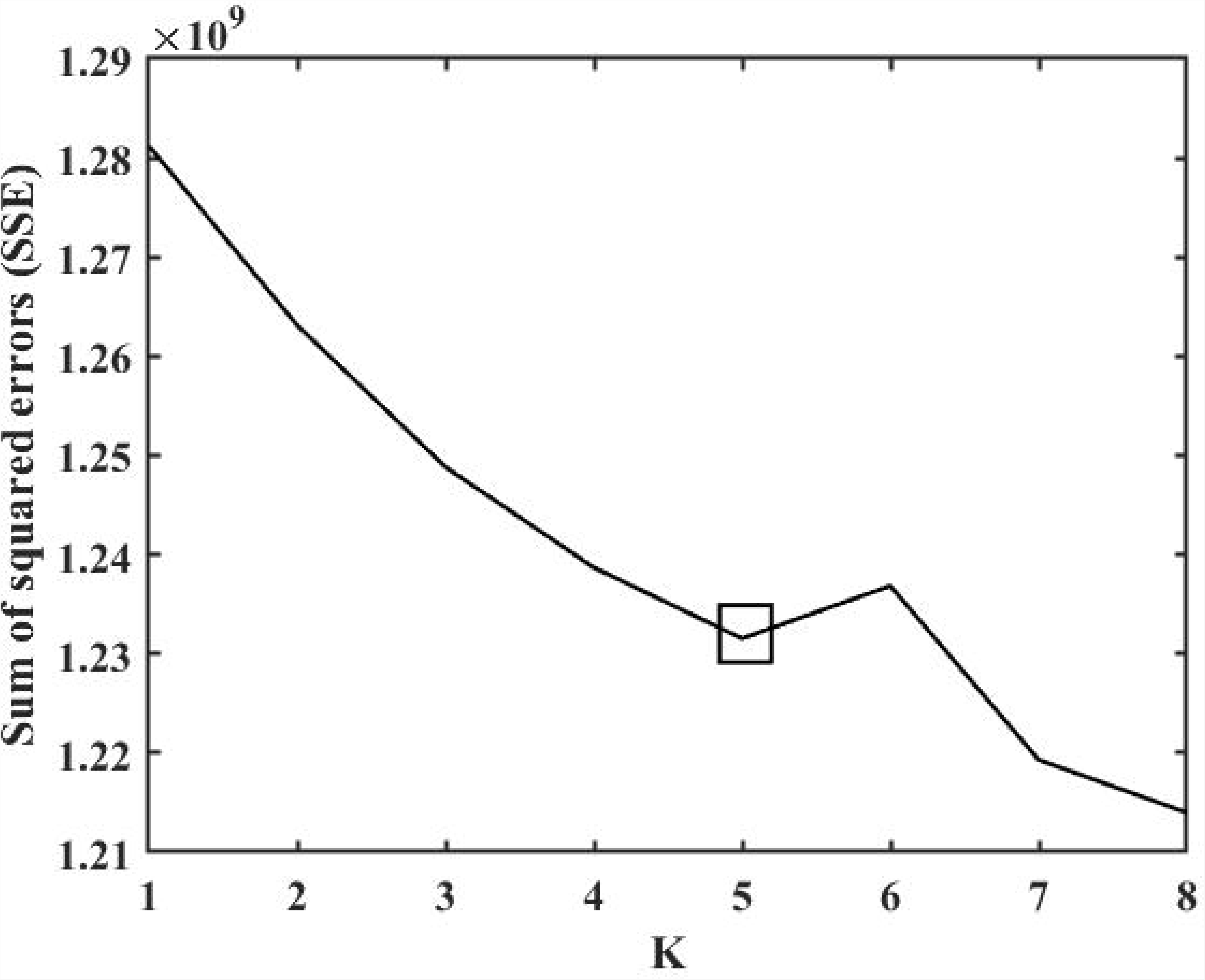

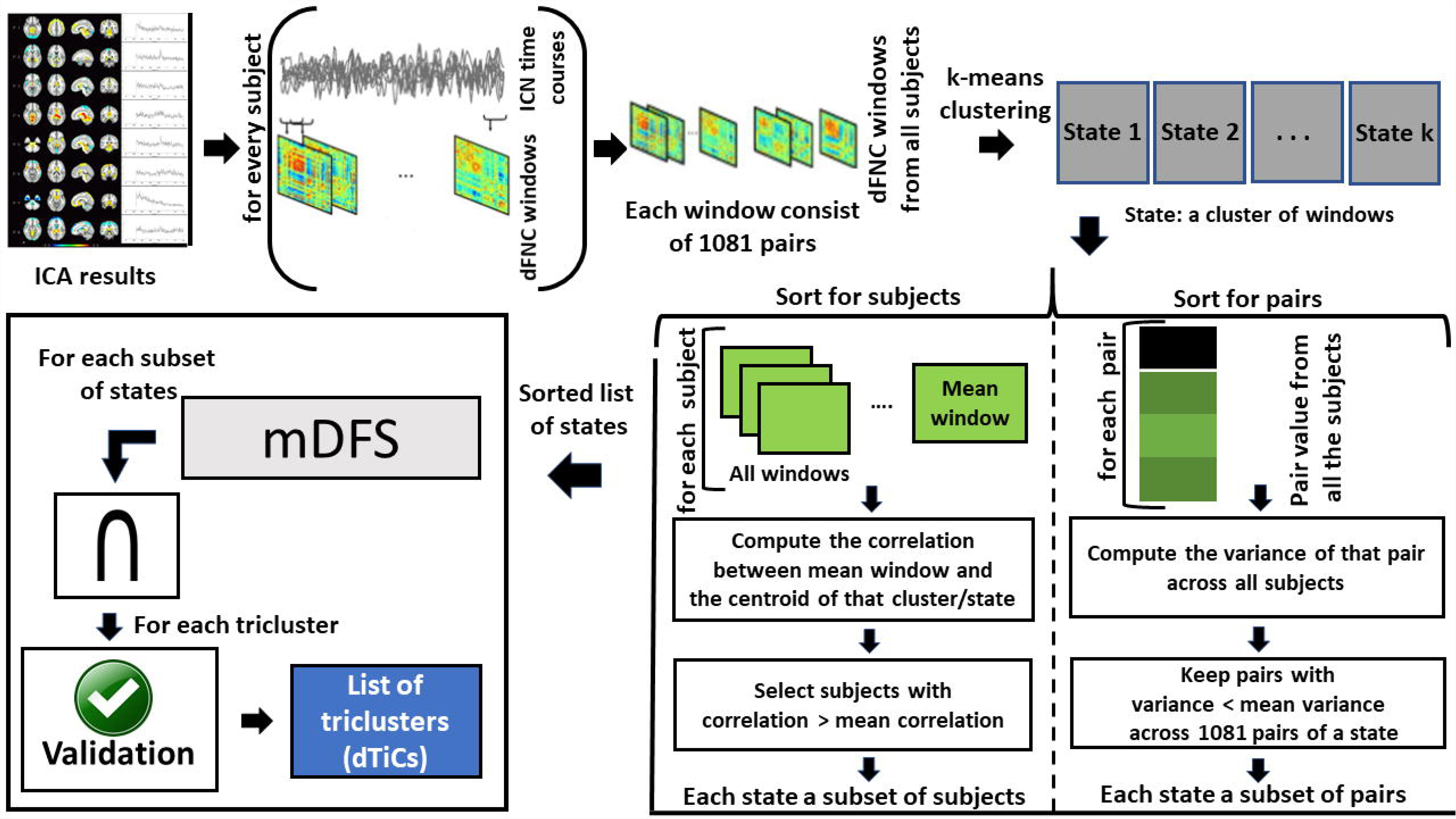

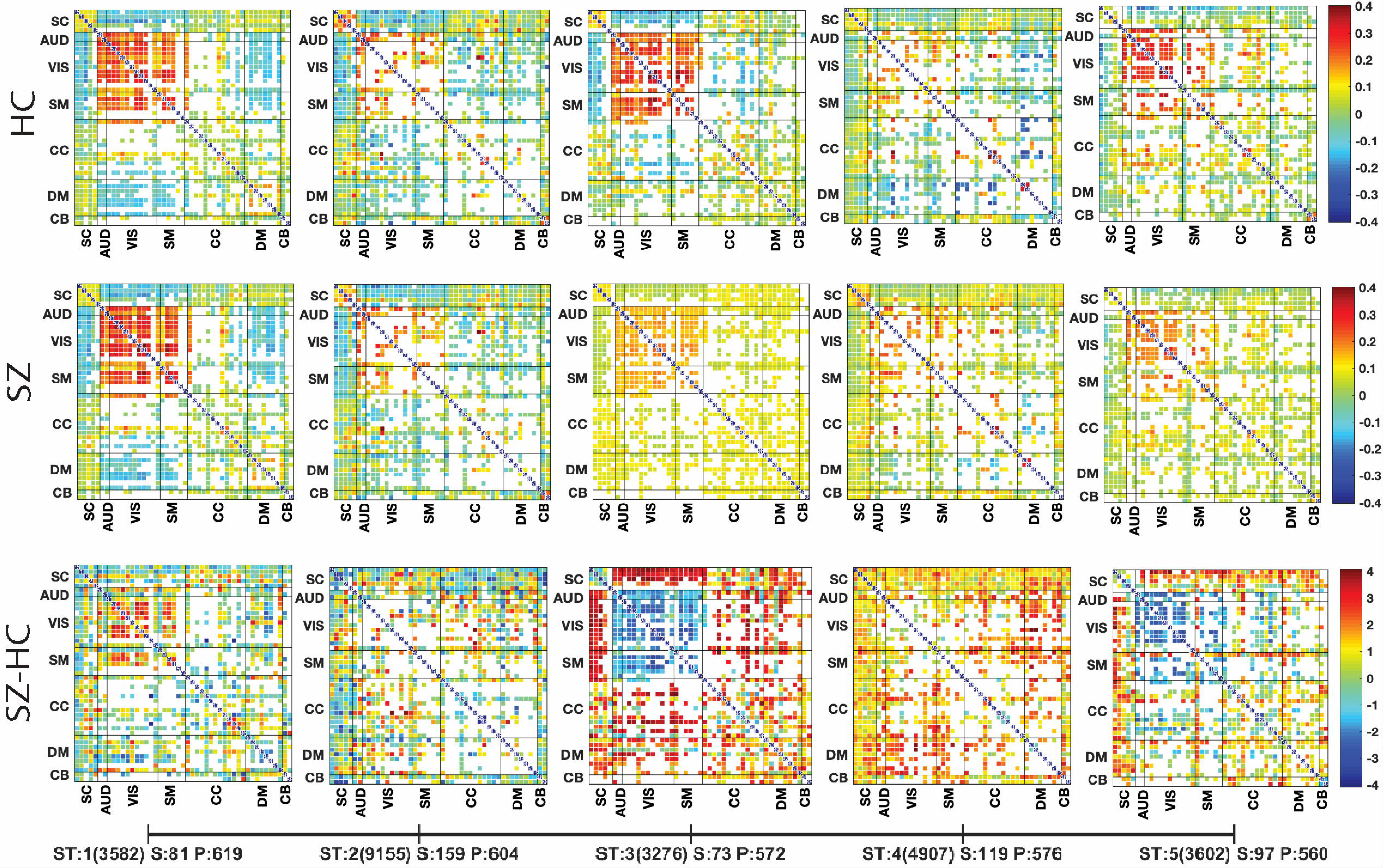

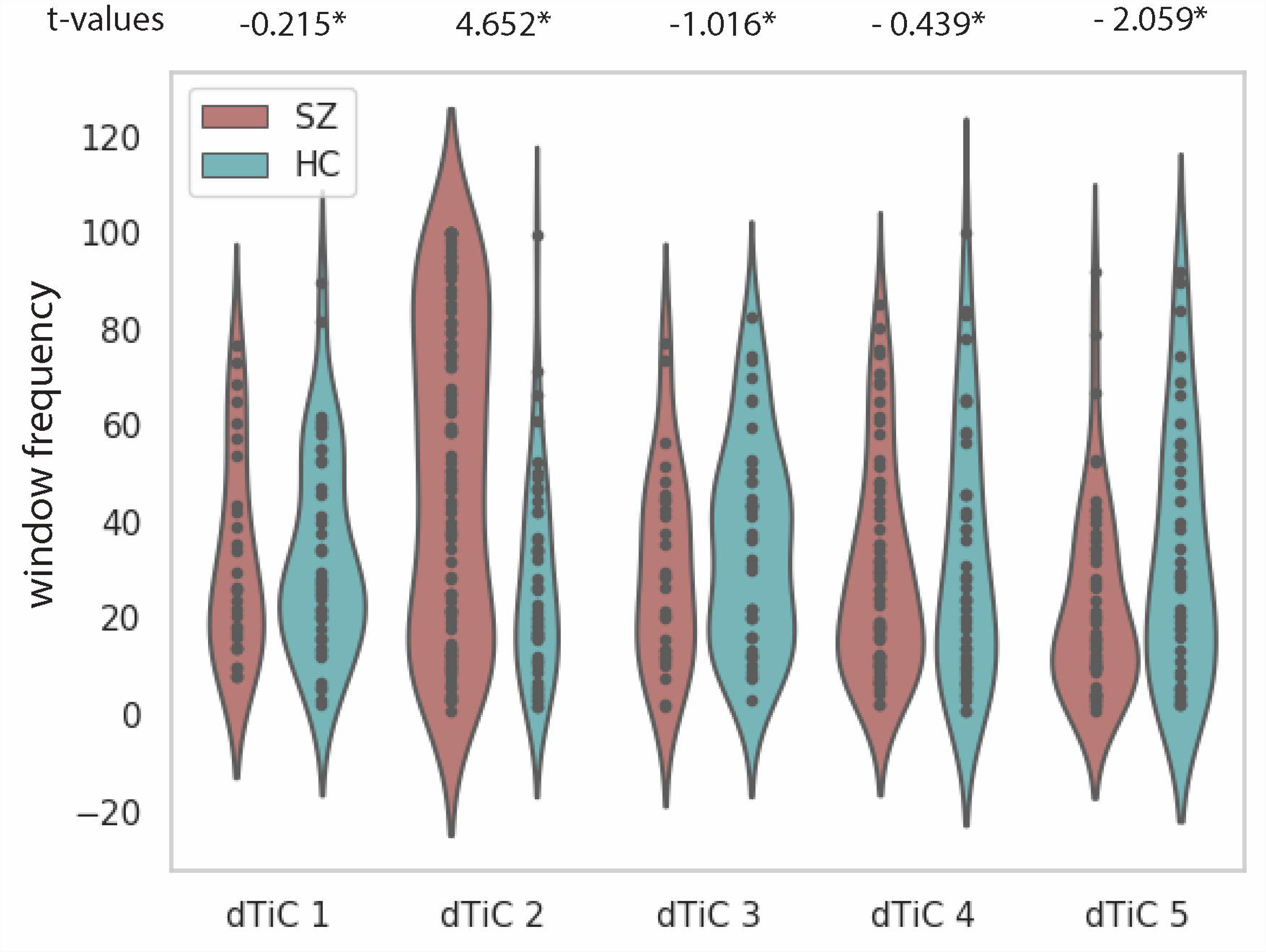

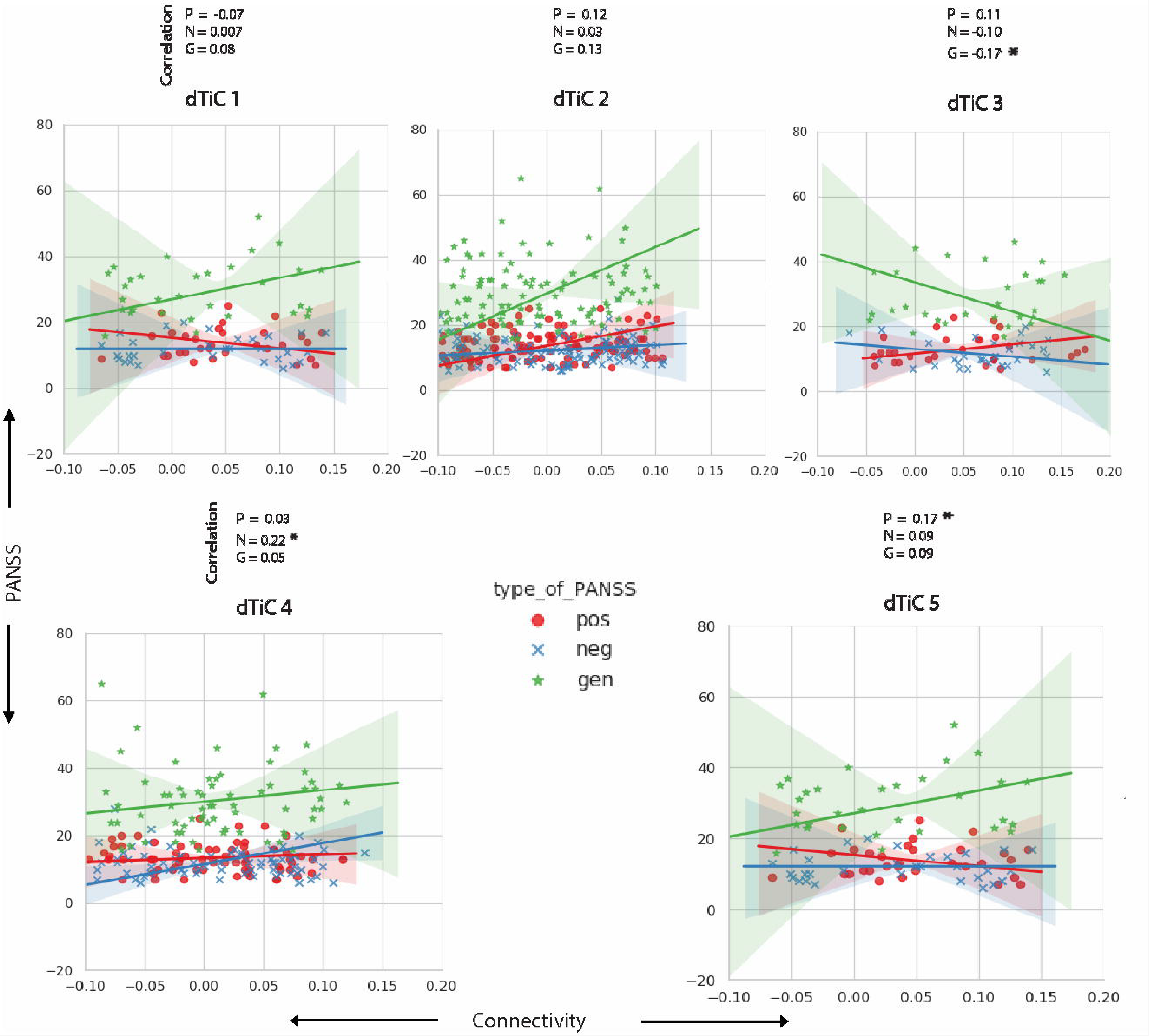

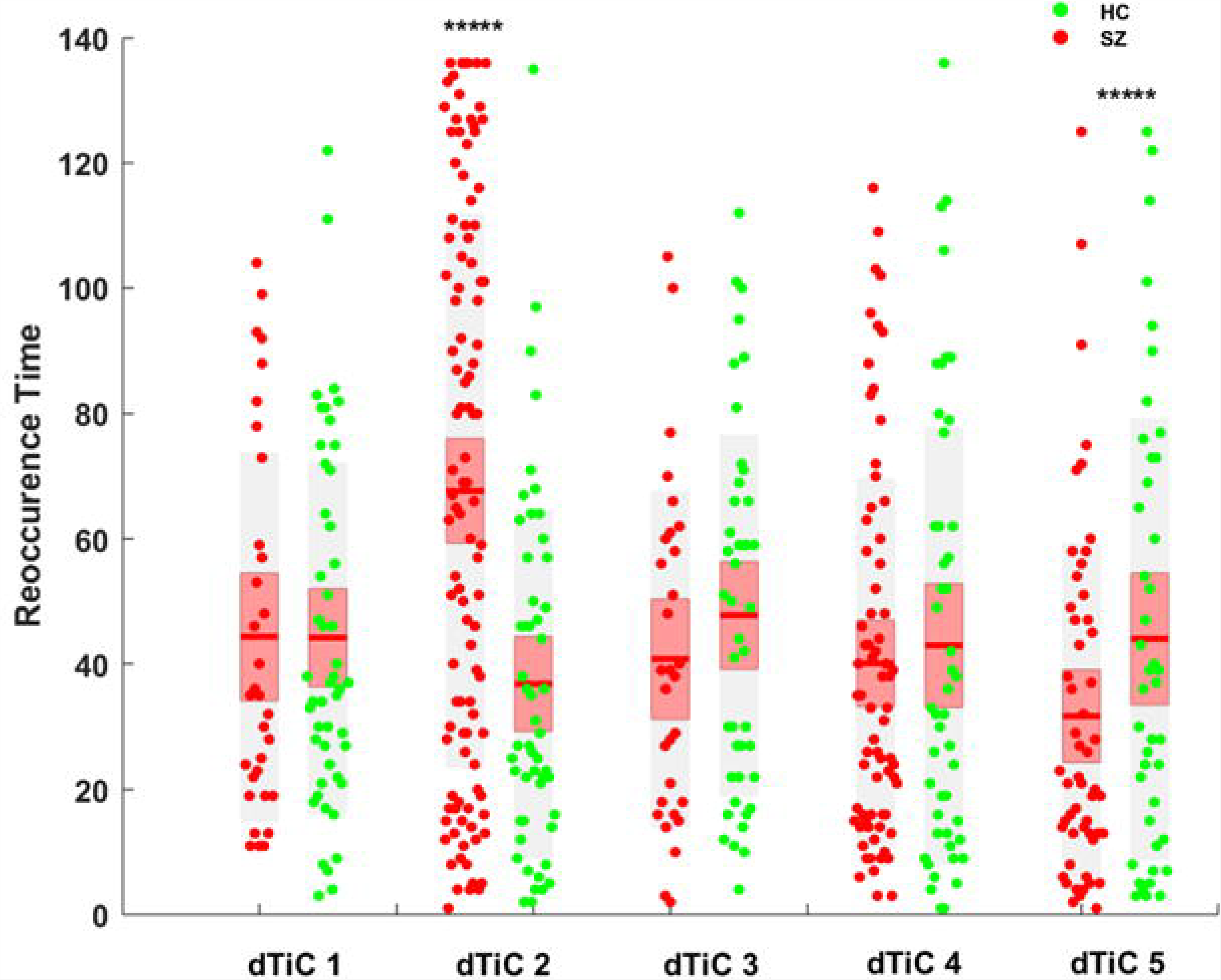

## Footnotes

1 http://afni.nimh.nih.gov/.

2 http://www.fil.ion.ucl.ac.uk/spm/.

3 http://trendscenter.org/software/gift.

