## Supplementary Information for "A novel method for tri-clustering dynamic functional network connectivity (dFNC) identifies significant schizophrenia effects across multiple states in distinct subgroups of individuals"

**dNTiC for a higher K (K = 9)**

We run the experiment with a higher K for checking the stability and changes in dTiCs extracted by our algorithm. After running k-means for K = 9 we run dNTiC with the identical settings of run 1 (M =1) and got the following tri-clusters presented in figure 1.

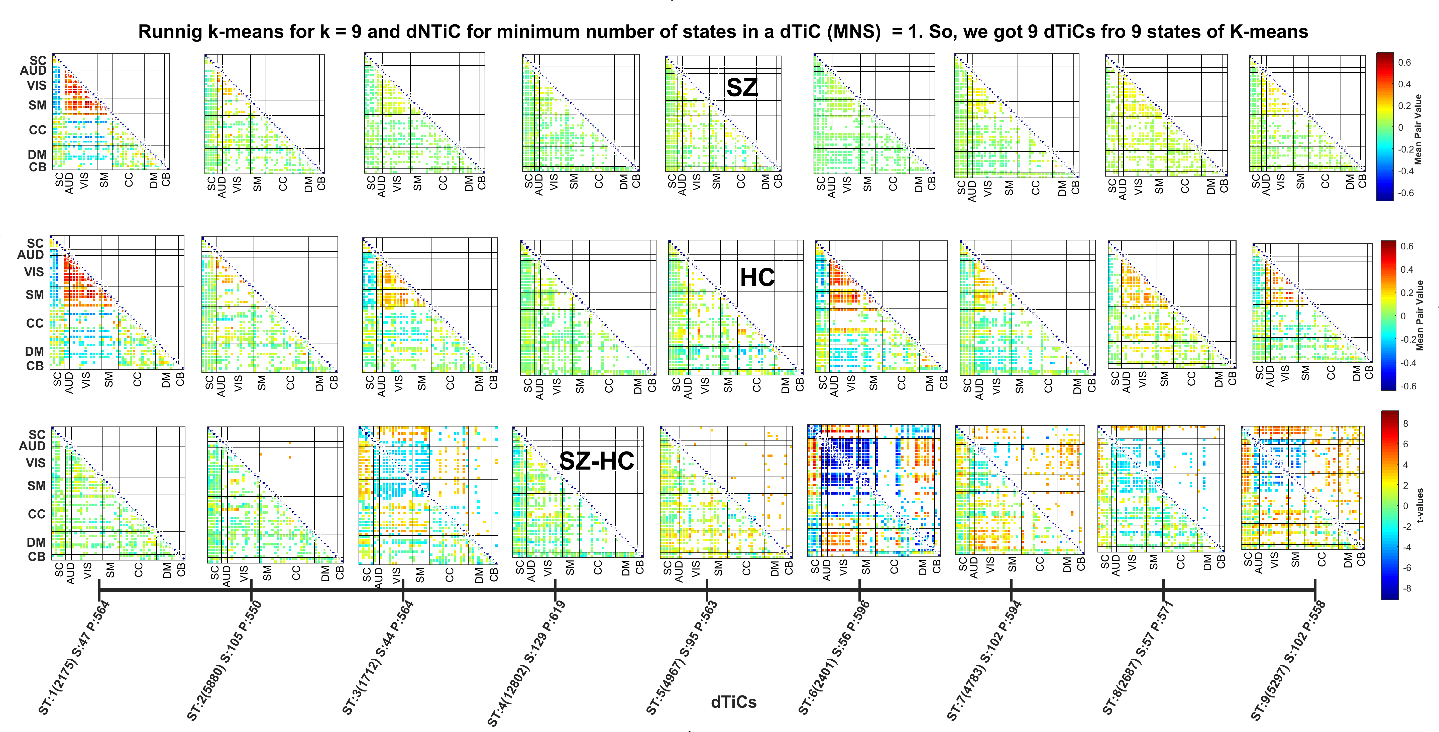

**Fig 1.** Schizophrenia (SZ) – Healthy control (HC) t-values for k = 9 and the minimum number of required window type in a tri-cluster (dTiC) is set to 1. Tri-cluster 2 3, 5, 6, 7, 8, 9 passed the correction. The upper triangle of the figures in the bottom row indicates the false discovery rate (FDR) corrected pairs.

We can see dTiC 3, 6, 8 and, 9 show major group differences. From the results, we can observe all the tri-clusters from run 1 have been replicated in these dTiCs. Figure 1 show group differences in SC, VIS, and SM domain across the tri-clusters. dTiC 3 shows differences in DM to all other domains and the direction of the difference is SZ > HC. dTiC 1, 2, and 4 showed no significant group differences. Apparently, results we presented in figure 1 replicates all information from earlier run (k = 5) where dTiC 6 is the replication of dTiC 3 of figure 1.

However, we extend the method for extracting more fine-grain subgroups of the dataset consequently, observing group differences in a more precise manner. We run the method for an increased number of states in is tri-cluster which is theoretically mounting the connectivity pattern to check their persistence across the time course. Multiple states capture more longer connectivity dynamics. We described the prolonged analysis in the following paragraphs.

**Run 2 (N = *, M = 2 and P = *; * stands for wild card)**

For run 2, we increased the input parameter k from 1 to 2. Now, dNTiC extracts tri-cluster which includes at least 2 states. By increasing M, we investigate the homogeneity to a bigger extent, across multiple states. From figure 2, we can see a different subset of states has been included in every dTiC and we got a total 4 dTiCs for this run. Most of the clusters show the almost identical contribution from both groups (SZ/HC) in each pair of dTiC 1 and 2. However, there are few significant cells that are scattered across those dTiCs, mostly in DM and CB for dTiC 1 and CC for dTiC 2. dTiC 1 has both directionalities HC > SZ (blue) and SZ > HC (red) where dTiC 2 has only SZ > HC (only reddish). dTiC 3 shows significant connectivity in VIS, SM (SZ > HC) and few HC > SZ in SC to other domains. dTiC 4 shows mostly HC > SZ group differences across all over the dTiC. In the earlier run (figure 3) dTiC 3 and 4 showed maximum significant group differences. In this run, dTiC 2, 3 which includes state 2, 3 does not survive the FDR correction.

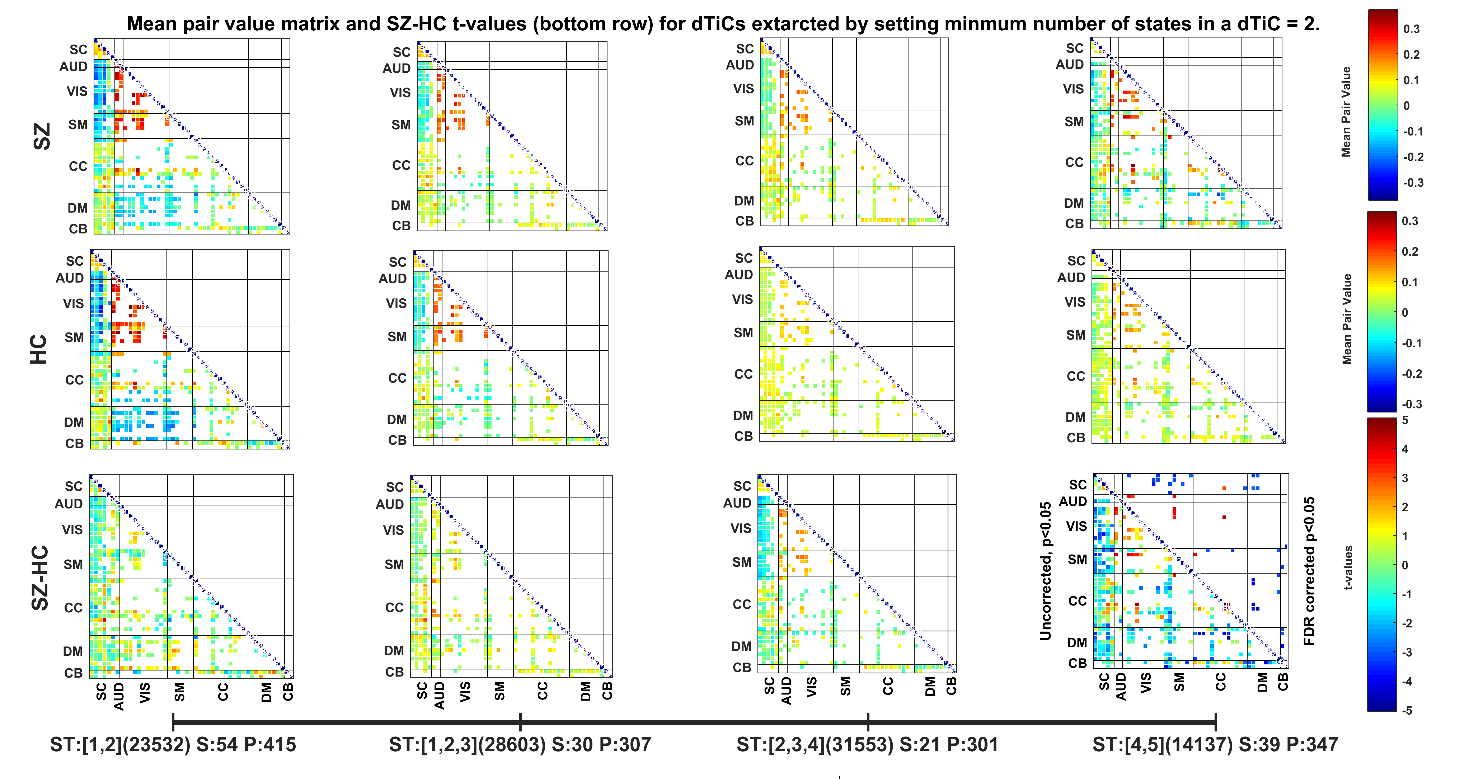

**Fig 2.** Schizophrenia (SZ) – Healthy control (HC) t-values for 4 tri-clusters (dTiCs) when we set the minimum number of required states in a dTiC is equal to 2. The algorithms captured inter window-clusters relation across different subsets of 5 window states. It evaluated intersection across the pairs and subjects of different states included in a dTiC. We can see the difference in the number of pairs/subjects from the figure. Xticklabel is consistent with Fig1, except we observed multiple states (window cluster) included in all the dTiCs.

The only significant tri-cluster is dTiC 4 which clustered states 4 and 5 together. The number of subjects is 39 and pairs are 347 which is approximately half of any dTiC from the earlier run. So, it is sort of separate subgrouping with distinct connectivity patterns which resulting in the group differences. The significant group differences are SC to VIS, SM, DM regions in this tri-cluster.

Figure 3 demonstrates the state analysis results from previous study (Damaraju et al., 2014) for a quantitative comparison with our proposed method.

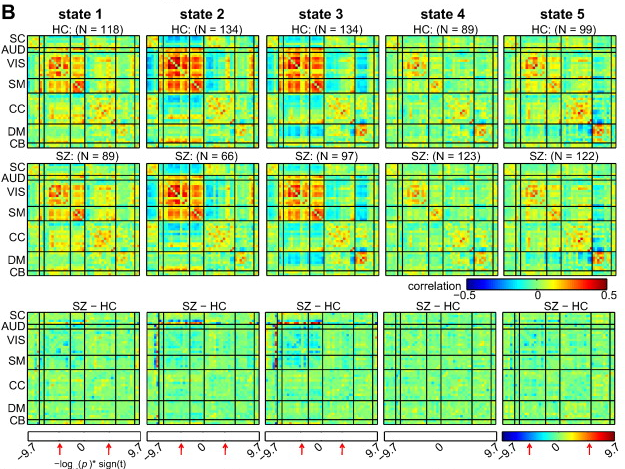

**Fig 3.** SZ – HC group differences from the previous study (Damaraju et al., 2014). The states extracted by k-means on dFNC time course. We observe less group differences comparing to our dTiCs (Fig. 4 in main text) in all clusters. Also, the states look very similar for SZ and HC subjects. A better clustering and sorting of the states yield more group differences across multiple brain regions.
