## Supplementary material for "A novel method for tri-clustering dynamic functional network connectivity (dFNC) identifies significant schizophrenia effects across multiple states in distinct subgroups of individuals": Figure Captions

**Figure legend/captions**

**Figure 1.** Dynamic functional network connectivity (dFNC) dataset. Different color represents a distinct window, and each window has 1081 (u_1_ to u_1081_) values where the value represents the strength of connectivity between a pair of components.

**Figure 2**. The elbow criteria for determining the model order (k) for k-means. We run k-means for k =1 to 8. Y-axis represents the sum of squared errors (SSE) for each k in the X-axis. The rectangular box indicates the elbow at k =5.

**Figure 3.** Our proposed tri-clustering methodology. It can be divided into three basic steps, the first one where it clusters the windows from all the subjects into a certain number of clusters/states using standard k-means. Then, the method starts sorting the states in two dimensions (subject, pair) and describes each of the states as a subset of subjects and a subset of pairs. Finally, the sorted states go through a homogeneity maximization procedure toward forming the tri-clusters by exploring all possible subsets of a given set of states using a modified depth-first search (mDFS) technique.

**Figure 4.** Group means (top two rows) and group differences (bottom row) of functional connectivity (FC) in distinct tri-clusters (dTiC). Schizophrenia (SZ )and healthy control (HC) individuals are included in each dTiC. We evaluated the group-wise contribution on each pair of a dTiC by both groups. Two sample t-tests using a null hypothesis of “No group difference” we used to compare the mean of patients vs. the mean of controls. A higher t-value indicates the rejection of the null hypothesis irrespective of their sign. However, the sign of t-values represents the directionality of the group difference. The pair matrix (47 x 47) is labeled into 7 different brain domains subcortical, auditory, visual, sensorimotor, cognitive control, default-mode, cerebellar, respectively. The white cells in the matrix indicate either the absence of that pair or non-significant group differences. The Xticklabel represents tri-cluster states (ST); the number of windows (of type ST) in that cluster from all the subjects included S: number of subjects in this dTiC; P: number of pairs included. These are the false discovery rate (FDR )corrected differences.

**Figure 5.** Schizophrenia (SZ) – Healthy control (HC) t-values from a two-sample t-test computed on window frequency in each subject within the tri-cluster. Window frequency is defined as the percentage of a specific type of window in a subject. The violins demonstrate the distributions of the data points. The asterisk (*) sign on the t-value indicates the statistical significance at a level of *p* < 0.05.

**Figure 6.** Correlation between positive and negative syndrome scale (PANSS) score and mean connectivity strength for schizophrenia (SZ) subjects within each subgroup (dTiC). The subplots represent the data points and the regression lines between the variables. We plotted mean connectivity (X-axis) vs. PANSS score (Y-axis). In subgroup 1, the SZ subjects are anti-correlated with positive symptoms - the positive PANSS scores decrease with the rise of connectivity strength. Mentionable, these subjects show higher connectivity in visual (VIS) and sensorimotor (SM) regions (figure 5). Subjects in dTiC 3, 5 are highly correlated with positive symptoms and exhibit lower connectivity strength than controls within these subgroups. Patients in dTiC 3 show slightly diminishing trends in negative PANSS scores with the increase of their connectivity level. Anti-correlation with negative PANSS scores might indicate functional dysconnectivity in auditory (AUD), VIS, and SM regions (figure 5) effect negative symptoms in schizophrenia. dTiC 4 has a strong association with negative symptom scores and show very sparse connectivity in VIS and SM regions (figure 5). The significant correlations are indicated using the asterisk (*) sign.

**Figure 7.** Reoccurrence time of both SZ and HC subjects included in each dTiC. The green and red circles represent the reoccurrence time of HC and SZ subjects, respectively. The red line on the box shows the population mean—the asterisk (*) sign indicates the dTiC with statistically significant differences between SZ and HC groups. We run a two-sample t-test on the reoccurrence time of SZ and HC subjects within a tri-cluster to evaluate the group differences. Subjects in dTiC 2 and 5 show statistical significance at a level of *p* < 0.05.

**Supplementary Figure 1.** Schizophrenia (SZ) – Healthy control (HC) t-values for k = 9 and the minimum number of required window type in a tri-cluster (dTiC) is set to 1. Tri-cluster 2 3, 5, 6, 7, 8, 9 passed the correction. The upper triangle of the figures in the bottom row indicates the false discovery rate (FDR) corrected pairs.

**Supplementary Figure 2.** Schizophrenia (SZ) – Healthy control (HC) t-values for 4 tri-clusters (dTiCs) when we set the minimum number of required states in a dTiC is equal to 2. The algorithms captured inter window-clusters relation across different subsets of 5 window states. It evaluated intersection across the pairs and subjects of different states included in a dTiC. We can see the difference in the number of pairs/subjects from the figure. Xticklabel is consistent with Fig1, except we observed multiple states (window cluster) included in all the dTiCs.

**Supplementary Figure 3.** SZ – HC group differences from the previous study (Damaraju et al., 2014). The states extracted by k-means on dFNC time course. We observe less group differences comparing to our dTiCs (Fig. 4 in main text) in all clusters. Also, the states look very similar for SZ and HC subjects. A better clustering and sorting of the states yield more group differences across multiple brain regions.
